# TidyGEO: Preparing analysis-ready datasets from Gene Expression Omnibus

**DOI:** 10.1101/2023.02.09.527930

**Authors:** Avery Mecham, Ashlie Stephenson, Badi I. Quinteros, Grace Salmons, Stephen R. Piccolo

## Abstract

TidyGEO is a Web-based tool for downloading, tidying, and reformatting data series from Gene Expression Omnibus (GEO). As a freely accessible repository with data from over 4 million biological samples across more than 4,000 organisms, GEO provides diverse opportunities for secondary research. Transcriptomic data are most common in GEO, but other measurement types are also prevalent, including DNA methylation levels, genotypes, and chromatin-accessibility profiles. GEO’s diversity and expansiveness present opportunities and challenges. Although scientists may find assay data relevant to a given research question, most analyses require sample annotations, such as a sample’s treatment group, disease subtype, or age. In GEO, such annotations are stored alongside assay data in delimited, text-based files. However, the structure and semantics of the annotations vary widely from one series to another, and many annotations are not useful for analysis purposes. Thus, every GEO series must be tidied before it can be analyzed. Manual approaches may be used, but these are error prone and take time away from other research tasks. Custom computer scripts can be written, but many scientists lack the computational expertise to create such scripts. To address these challenges, we created TidyGEO, which supports essential data-cleaning tasks for sample-level annotations, such as selecting informative columns, renaming columns, splitting or merging columns, standardizing data values, and filtering samples. Additionally, users can integrate annotations with assay data, restructure assay data, and generate code that enables others to reproduce these steps. The source code for TidyGEO is at https://github.com/srp33/TidyGEO.

## Background

Gene Expression Omnibus (GEO) is an Internet-based, publicly accessible repository for high-throughput molecular-abundance data^1,2^. Originally, GEO was designed for gene-expression profiles^3^, but its scope has broadened, now housing data for other measurement types, including DNA methylation levels, genotypes, and chromatin-accessibility measurements. GEO data are stored in four units: *Platforms*, which define a set of molecules that may be detected using a given profiling technology; *Samples*, which describe measurements produced for a single replicate; *Series*, which organize assay data and metadata for a set of samples that make up an experiment; and *Datasets*, which are curated collections of samples for a particular platform^2^. Most samples are not part of a Dataset due to the time and expertise required for curation.

GEO enables researchers to share data with the broader community. Many journals and funding agencies require such sharing to enable validation of research results and to ensure that data assets are accessible^4^. GEO data are used for many types of secondary research. Some researchers use GEO data for methods development and evaluation. For example, Zhou, et al. combined diverse datasets to impute gene-expression values with the goal of maximizing platform compatibility^5^; Eren, et al. used GEO data to compare the effectiveness of biclustering algorithms^6^; and Golightly, et al. curated a compendium of GEO datasets to enable benchmark comparisons across machine-learning algorithms^7^. Other researchers have used GEO data for discovery, such as identifying differentially expressed genes^8,9^, identifying pathways that influence disease development^10,11^, or investigating the potential to repurpose existing drugs^12^.

GEO adheres to the Minimum Information about Microarray Experiments (MIAME) and Minimum Information about a high-throughput SeQuencing Experiment (MINSEQE) guidelines, which define the types of content that should be provided in publicly available, gene-expression datasets^13,14^. This content can be categorized as 1) metadata about the experiment, 2) sample-level annotations, and 3) a processed version of the molecular-assay data. Metadata include information such as a title for the study, the species name(s), an experimental design description, contact information, and the platform(s) used. Sample-level annotations typically indicate the experimental condition(s) associated with each research subject and covariate factors such as the age, sex, and/or disease subtype of each subject; these annotations differ considerably across studies. Additional sample-level annotations are provided for informational purposes but may not be useful in analyses. Such variables might indicate the molecule type that was profiled, the extraction and hybridization protocols used, a description of how the data were preprocessed, the last-updated date, and subject identifiers used originally by the submitter. Although many GEO series provide a raw version of the data, many more provide a processed version of the data that were used in the researchers’ analysis. Examples of processed data include normalized microarray measurements, feature counts from read-aligned RNA-Sequencing data, RT-PCR measurements, etc.

GEO has long used a spreadsheet-based submission system to collect metadata and annotations from researchers. A curator reviews each submission. This process ensures that the information is structured consistently across studies and that key data elements are provided, while ensuring flexibility for a wide range of experiments. This flexibility has enabled GEO to grow quickly over the past decades and to meet its creators’ goal of remaining “flexible and responsive to future trends, rather than setting rigid requirements and standards for entry”^3^. However, challenges for data reuse^4^ have accompanied this flexibility. Within the constraints of the submission process, researchers provide free-form descriptions of sample characteristics. In some cases, researchers provide multiple values per cell, separated by delimiters. For example, a value of “female;52;anastrozole” might be used to represent a female breast-cancer patient who was 52 years old and had been treated with anastrozole. When reusing the data, a secondary researcher would need to decipher the semantic meaning of these values and write custom code to separate the values into distinct columns. In other cases, sample characteristics are stored as key-value pairs. For example, the same patient might be represented as “sex=female;age=52;drug=anastrozole”. This approach provides some semantic information, but secondary researchers would still need to parse the data points. In addition, missing values can cause problems. In some GEO datasets, sample-level annotations in a given row are shifted leftward to fill empty cells. Consequently, a given column may contain data for multiple variables. To analyze such data, a researcher would need to realign the values.

In many cases, these inconsistencies violate the “tidy data” principles, which state that every column should describe a single variable, every row should represent a particular observation (sample), and every table should represent a particular type of observational unit^15^. When data conform to these principles, they are conducive to diverse types of quantitative analyses, enabling researchers to devise analytical strategies that generalize across datasets.

Other challenges relate to semantics^16^. The names of columns often do not reflect the data stored in those columns. For example, Huang, et al. generated gene-expression data for Wilms tumor patients (GEO accession: GSE10320)^17^. Sample-level annotations indicate the clinical outcome for each patient: “Relapse” or “Non-relapse”. The column containing these values is labeled “characteristics_ch1”. As researchers analyze such data, they may wish to rename columns to be more descriptive. Additionally, the actual data values may lack standardization. For example, a more recent dataset published by the same group used “Yes” or “No” values to indicate whether a given Wilms tumor patient had relapsed^18^. If a secondary researcher wished to merge these datasets, they would need to modify the data to use a common vocabulary for describing relapse status. Finally, in some secondary analyses, researchers wish to use only a subset of the available samples. For example, they may wish to focus on Wilms tumor patients who had relapsed.

Aside from sample-level annotations, assay data often must be re-summarized. For example, Affymetrix microarray data are often summarized at the probeset level, and a given gene is often associated with multiple probesets. Researchers who wish to make gene-level inferences might wish to calculate the average (or median or maximum) value across all probesets for each gene.

Writing custom code to address any one of these problems may be trivial for some researchers. However, multiple such transformations are needed for many datasets, and these tasks differ widely among datasets^7^. Performing these tasks idiosyncratically for each dataset is inefficient and makes secondary analyses less accessible to researchers who lack computational skills. To address this problem, we created *TidyGEO*, an interactive Web application that enables researchers to download, tidy, and restructure GEO series into a form that is suitable for downstream analysis. Unlike alternative tools that focus primarily on molecular-assay data, TidyGEO emphasizes tidying sample-level annotations. However, TidyGEO also provides options for tidying molecular-assay data. Users can create graphical summaries of the data and download graphics files. Additionally, users can export the data in diverse formats.

In this article, we describe TidyGEO’s functionality in more detail and describe our findings from using it to tidy 109 GEO series. Furthermore, we highlight the importance of computational reproducibility and describe TidyGEO’s approach to ensuring that the tidying steps can be reproduced. Finally, we describe the landscape of existing tools for finding, interoperating, and reusing GEO data^4^; and we indicate how TidyGEO fits within that landscape.

## Results

TidyGEO enables researchers to tidy sample-level annotations, assay data, and platform annotations for GEO series. In addition, researchers can merge these data types, export the data in various formats, and generate scripts that support computational reproducibility. To evaluate our assertion that most GEO series need to be tidied and to verify that TidyGEO is effective for diverse types of GEO data, we used TidyGEO to import 109 series, identified needs for tidying each series, performed the tidying steps, and recorded which steps we performed. We selected these series in a semi-random manner with the intent to ensure a diverse representation of molecular-profiling platforms (Figure 1) and samples sizes. However, we emphasized larger sample sizes to test the limits of TidyGEO; the sample sizes ranged between 2 and 12,288 (Figure 2). We excluded datasets with sample sizes of zero or one because these were either incomplete or did not require help with formatting.

**Figure 1:**
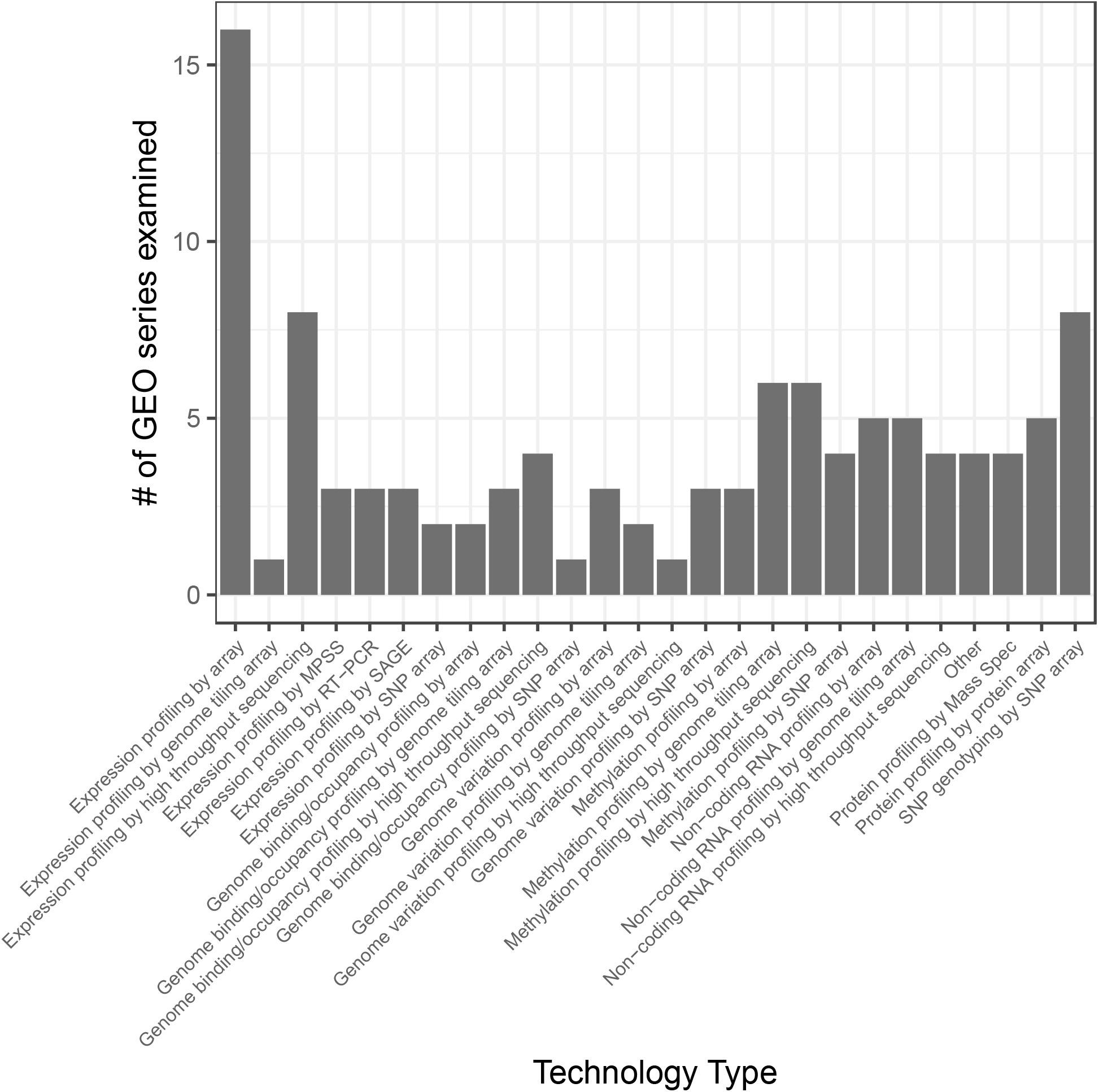
Assay types for the GEO series that we tidied to demonstrate TidyGEO’s functionality. We tidied 109 GEO series representing a diverse range of sizes and molecular-profiling technologies. This graph illustrates the number of datasets tested for each profiling technology.

**Figure 2:**
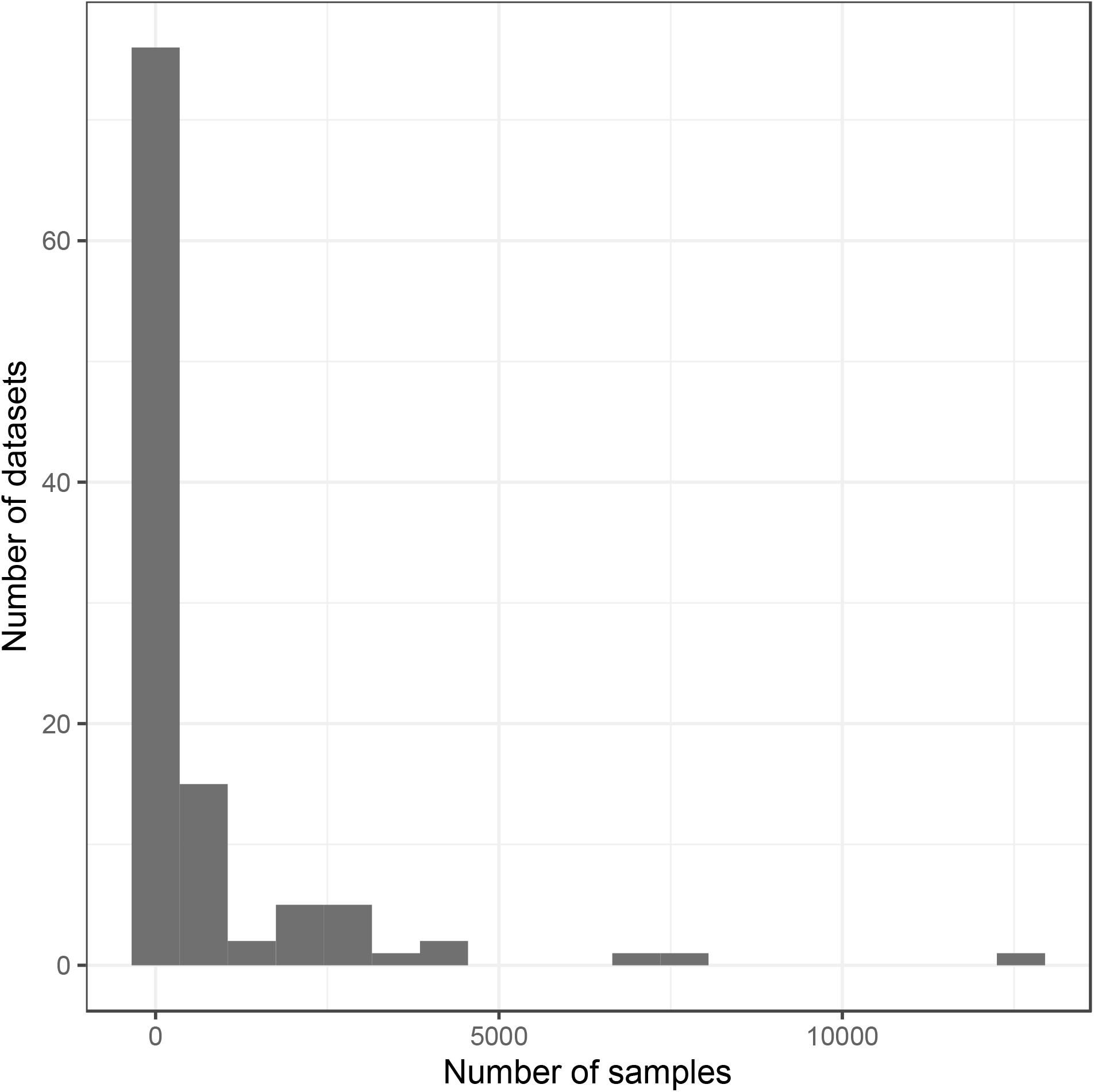
Sample sizes for the GEO series that we used to demonstrate TidyGEO functionality. We tidied 109 GEO series representing a wide range of sample sizes. This graph illustrates samples sizes across the series.

In TidyGEO, the “Clinical data” panel provides tabs for selecting informative columns, shifting cells, splitting key-value pairs, splitting columns, renaming columns, substituting values, and filtering samples. Table 1 lists the criteria we followed to assess whether each task needed to be performed for a given GEO series. We ignored the “Rename columns” and “Filter samples” tabs because these tasks are subjective and/or depend on the analysis being done. The “Select informative columns” tab provides preset filters that enable users to exclude columns that have web addresses (e.g., URLs for raw files), dates (e.g., upload dates), or for which all values are the same (e.g., tissue type). It also provides an option for the user to remove columns in a custom manner. We considered these two processes as separate types of tasks. If a series included multiple platforms, we used data for the first one listed.

**Table 1:** Criteria used to evaluate which tidying steps were needed for a given dataset. We used TidyGEO to process 109 datasets across a range of platforms and data sizes. In processing the data, we used these criteria to evaluate whether each type of tidying functionality needed to be applied to each dataset.

| Task | Criteria |
| --- | --- |
| Filter by column name | Does any column have data that you judge to be non-informative for data analyses? |
| Shift cells | In some cases, when values are missing, sample-level annotations become shifted leftward to fill empty cells. For many datasets, GEO has rectified this problem with new columns. Is there any cell in which this has occurred (and has not already been rectified)? |
| Split key-value pairs | Some sample-level annotations are represented as key-value pairs. In these cases, a data value is prefixed with a descriptor (key). Typically, the same descriptor is used for each cell. To analyze such data, it is helpful to use the descriptor as the column name and remove the descriptor from the cells in that column. Does any column have key-value pairs? |
| Split columns | In some cases, multiple values are stored in a single cell. For example, a person's age and sex might be stored in the same column, separated by commas. In these cases, it is helpful to split the column into multiple columns so the data can be analyzed more easily. Does any column need to be split? |
| Substitute values | Different researchers often use different terms to describe the same concepts. For example, one researcher might use "F" to indicate that a sample is female, while another researcher might use "Female" or "Woman." In other cases, spelling mistakes or inconsistencies might be present. Or delimiters might be used inconsistently. Standardizing those values can make it easier to analyze the data and combine datasets. Do you judge that any of the informative columns have values that should be standardized (substituted with different values)? |

Figure 3 illustrates the number of tidying tasks per series. All 109 series required at least two tidying tasks. Three series required all six tasks. Figure 4 indicates the frequency with which each task was performed. By far, the most common tasks involved selecting informative columns. But each of the six tasks was necessary for at least 10% of the series.

**Figure 3:**
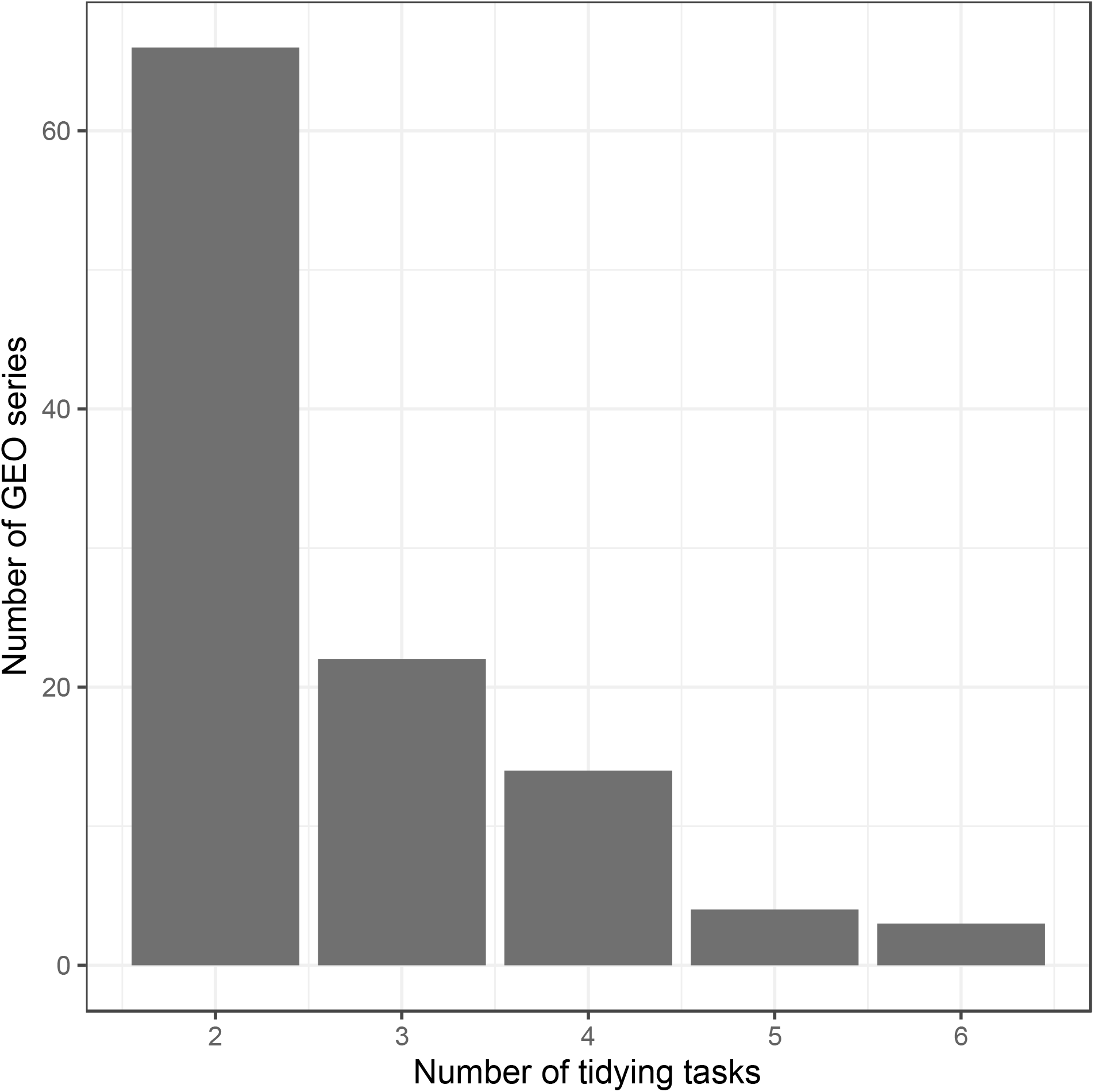
Number of tidying tasks necessary to tidy each of 109 GEO series. We used TidyGEO to tidy sample-level annotations for 109 GEO series. This graph illustrates the total number of tasks per series that were necessary to tidy the data. Two tasks were possible for the “Select columns” tab, and we did not consider the “Rename columns” or “Filter samples” tabs. So the maximum possible tasks per series was 6.

**Figure 4:**
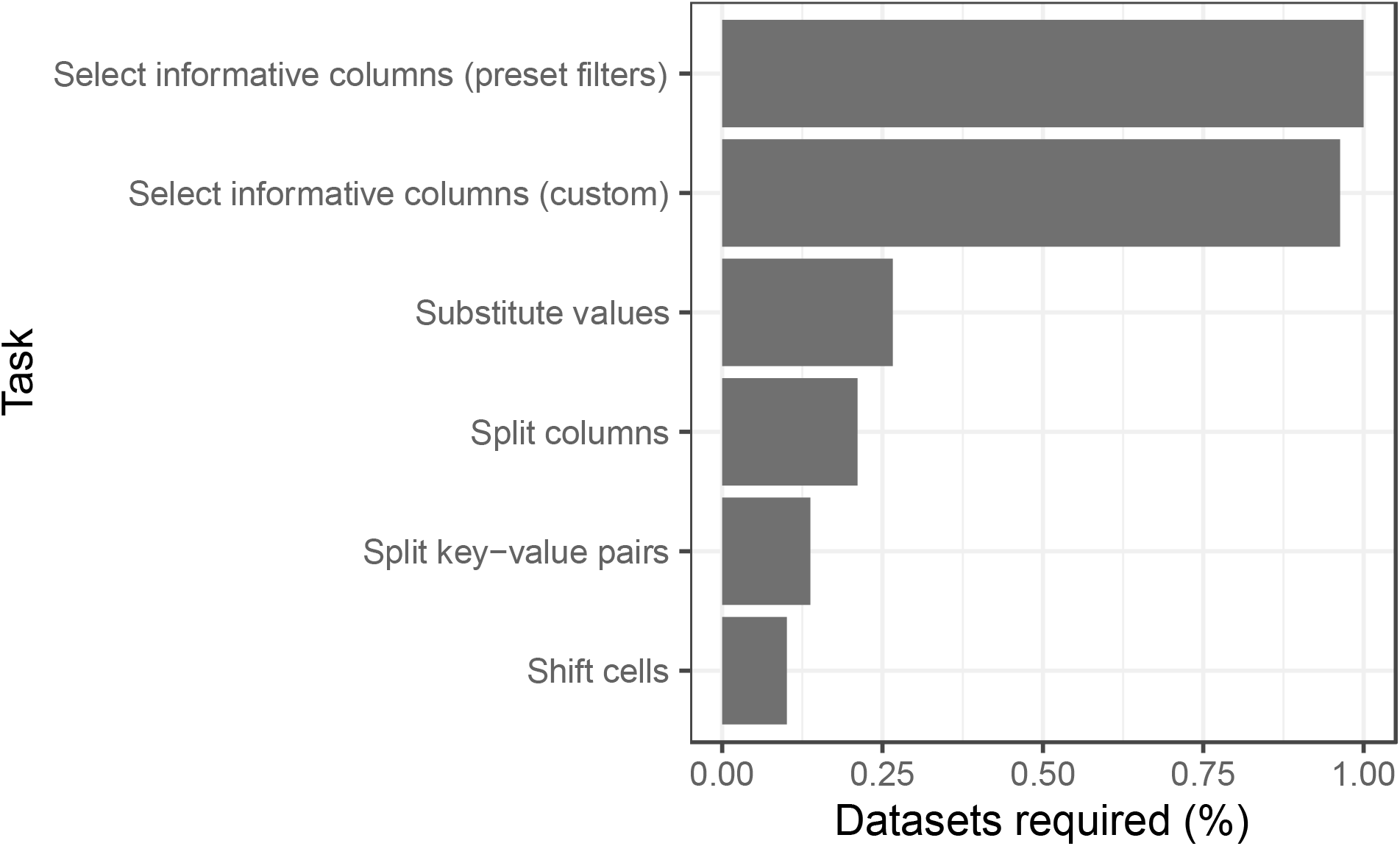
Percentage of times that each tidying task was needed to tidy each of 109 GEO series. We used TidyGEO to tidy sample-level annotations for 109 GEO series. This graph illustrates the percentage of times that each type of tidying task was performed across all series.

During testing, we found that assay data were not available for direct download for approximately 30% of datasets. In many cases (as in RNA-Sequencing experiments), raw data are stored in repositories separate from GEO (e.g., the Sequencing Read Archive^19^).

## Discussion

The vastness and accessibility of GEO data have inspired tools for finding, downloading, and parsing the data. Some of these tools are provided within GEO itself. From the GEO website, researchers can browse available data, perform differential-expression analyses^2^, and download data manually. To help in identifying relevant data, researchers can perform advanced searches. For example, they can filter based on metadata variables such as the title, authors, publication date, species name, or platform type of a given study. Furthermore, researchers can filter based on derived variables such as the sample count or availability of supplementary files. If desired, researchers can also perform such queries using an application programming interface. Other tools for working with GEO data have been created by third-party researchers. Below we describe many such tools and categorize them as a way to place TidyGEO’s functionality in context. In some cases, to provide historical context, we mention tools that are no longer available. In categorizing the tools, we refer to principles of making data findable, accessible, interoperable, and reusable (FAIR)^4^. Where applicable, we note gaps not currently filled by existing tools and how TidyGEO helps to fill those gaps.

### Finding GEO data

The first important step to reusing GEO data is to find the most relevant data for a given study. *GEOmetadb* is an R package that supports flexible querying of GEO metadata and thus may enable researchers to find relevant data more easily than using the GEO website^20^. *STARGEO* uses a crowdsourcing approach to annotate GEO series^21^; graduate students with a biomedical background have reviewed series metadata and manually assigned tags to them. Researchers can submit queries to identify series associated with topic(s) of interest. Another tool that used crowdsourcing is *CREEDS*; students in a massively open online course annotated GEO data and created themed collections of searchable gene signatures (lists of genes up- or down-regulated after a sample subset has been perturbed naturally or experimentally)^22^. OMics Compendia Commons (OMiCC) and *GEOMetaCuration* enable biomedical researchers to collaboratively annotate GEO data and then create and share cross-study compendia^23,24^. geoCancerPrognosticDatasetsRetriever is an R package that enables researchers to identify GEO datasets specific to cancer-prognosis research^25^. In an alternative approach, researchers have acted as curators. For example, *curatedOvarianData* is a compilation of expression data and metadata (mostly from GEO) associated with ovarian tumors^26^. Gemma has a much broader collection of curated gene-expression datasets^27^.

TidyGEO does not attempt to replicate these efforts. Instead, it asks researchers to search for data series using existing tools and then specify a series identifier (e.g., GSE10320) to begin the tidying process.

### Accessing GEO data

Via the GEO website, data are freely accessible using the HTTP/S and FTP protocols; each entry is assigned a unique identifier. Pointing and clicking can be cumbersome when researchers need to download many data series. As an alternative, scientists can use the GEOquery package to download GEO data programmatically^28^. TidyGEO uses this package to obtain data.

Tools like shinyGEO, Bioinformatics Array Research Tool, and GEOexplorer provide interfaces that allow researchers to search for GEO series and then retrieve those data^29–31^. Other repositories store subsets of GEO data; these include the curated packages mentioned above, as well as scanGEO^32^ and inSilicoDb (no longer available)^33^. Such repositories are typically limited to particular assay types (e.g., gene-expression microarray data).

GEO contains data for hundreds of thousands of human and mouse samples that have been profiled using RNA-Sequencing technologies. However, these data have been preprocessed using diverse reference genomes, gene annotations, and alignment/quantification tools. ARCHS4 uses cloud-based infrastructure to acquire the data in raw form (from the Sequence Read Archive^19^) and reprocess them so that researchers can more easily access and analyze the data^34,35^.

### Making GEO data interoperable

The FAIR guidelines recommend that data be compatible with tools available to process such data. GEO releases data in specific file formats that are consistent across series and platforms. However, these formats are specific to GEO, so most data-analysis tools are unable to import them directly. GEOquery transforms data into data frames and matrices so that analyses can be performed using the R statistical software^36^. *shinyGEO* uses GEOquery to obtain data and provides an option to download sample-level annotations in CSV format^29^. In addition, MeV and BRB-ArrayTools provide ways for researchers to import and export GEO datasets^37,38^. SMart Automatic Classification (SMAC) enables researchers to mine PubMed for datasets associated with user-defined queries, extract the data from GEO (using GEOquery), and then transform the data; it focuses primarily on cancer research, but some functionality can be applied more broadly^39^.

TidyGEO provides diverse options for exporting data, thus providing flexibility for researchers who wish to analyze data with external tools. In TidyGEO, data can be downloaded in any of five formats. In some cases, these formats are tied to specific software (e.g., Microsoft Excel or BRB-ArrayTools); in other cases (CSV, TSV, JSON), the formats are generic and thus can be imported into many tools for custom analyses.

### Reusing GEO data

In describing reuse, we focus on tools that provide *analytical* capabilities for gene-expression data. Researchers commonly seek to identify differentially-expressed genes^40^, perform functional-enrichment analyses, construct gene co-expression networks, identify disease subtypes via clustering, classify samples (e.g., predict survival, treatment response), perform meta-analyses (combine evidence across studies)^41^, and create visualizations. Many tools support such tasks(Table 2), typically via Web-based interfaces. Most of these tasks require sample-level annotations. For example, when performing a differential-expression analysis, researchers must place samples into comparison groups. Web-based analysis tools often extract metadata from GEO and ask the user to indicate the group associated with each sample. However, in many cases, it is useful to go beyond this capability. For example, if a researcher were comparing disease and normal samples, they might wish to account for race, sex, and/or age as covariates; we are unaware of existing tools that provide this flexibility. TidyGEO helps to fill the gaps in such scenarios. But rather than enabling researchers to analyze data directly, TidyGEO supports data-preparation tasks, enabling researchers to export analysis-ready data and import them into whatever third-party tool(s) are most suitable for the analysis tasks they wish to perform.

**Table 2:**
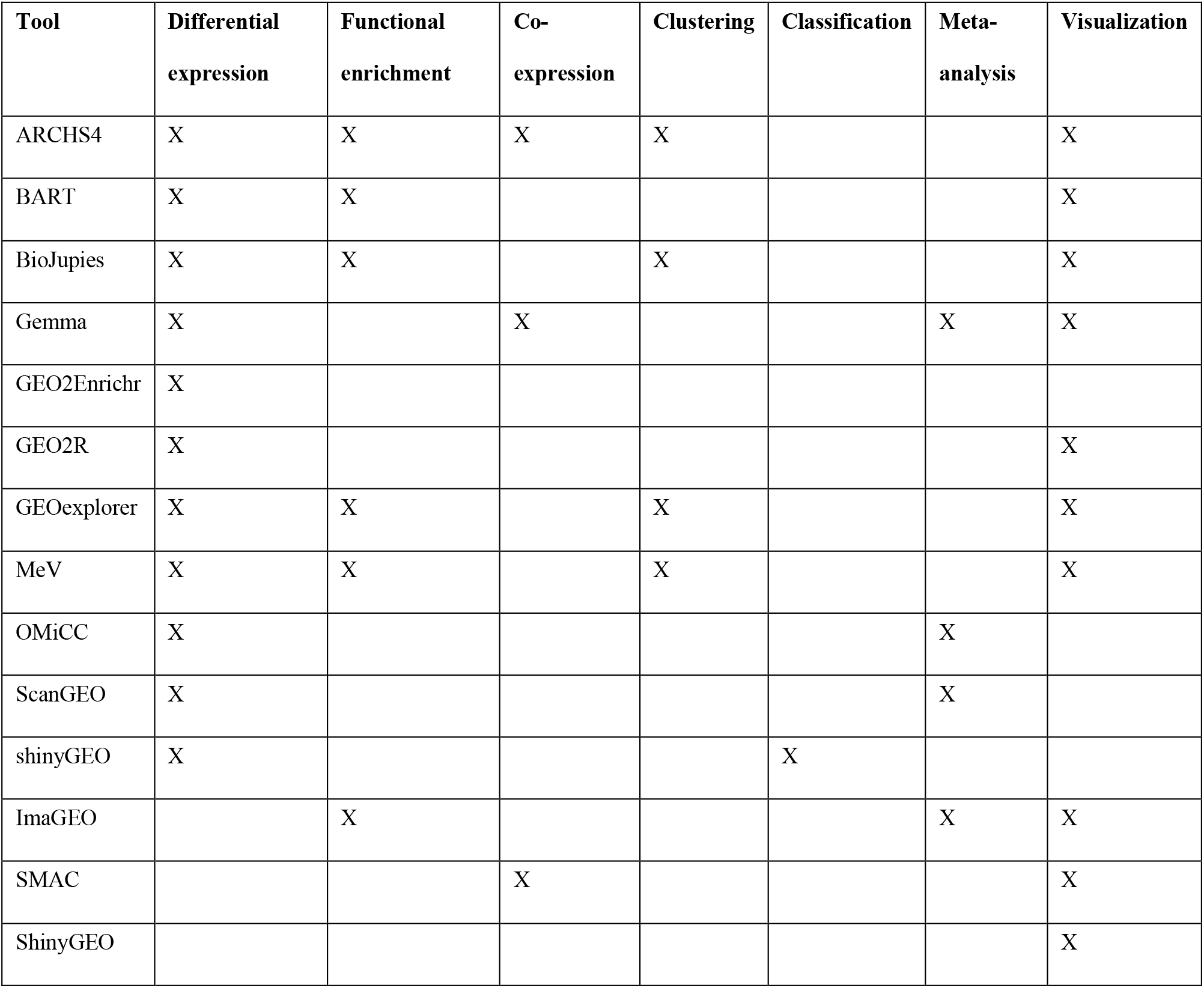
Summary of third-party tools that support reuse of GEO data.

| Tool | Differential expression | Functional enrichment | Co-expression | Clustering | Classification | Meta-analysis | Visualization |
| --- | --- | --- | --- | --- | --- | --- | --- |
| ARCHS4 | X | X | X | X |  |  | X |
| BART | X | X |  |  |  |  | X |
| BioJupies | X | X |  | X |  |  | X |
| Gemma | X |  | X |  |  | X | X |
| GEO2Enrichr | X |  |  |  |  |  |  |
| GEO2R | X |  |  |  |  |  | X |
| GEOexplorer | X | X |  | X |  |  | X |
| MeV | X | X |  | X |  |  | X |
| OMiCC | X |  |  |  |  | X |  |
| ScanGEO | X |  |  |  |  | X |  |
| shinyGEO | X |  |  |  | X |  |  |
| ImaGEO |  | X |  |  |  | X | X |
| SMAC |  |  | X |  |  |  | X |
| ShinyGEO |  |  |  |  |  |  | X |

## Conclusions

GEO data are largely findable and accessible. Furthermore, many tools are available to support interoperability and reusability of GEO data. However, a key gap that prevents researchers from taking full advantage of this resource is a lack of support for wrangling (tidying) the data into an analysis-read state. Data tidying requires human interaction and judgment from a domain expert. To be efficient and reproducible, it requires custom code. TidyGEO aims to meet these needs by facilitating the process of interacting with the data and generating custom code. Accordingly, we anticipate that it will be most useful to researchers who wish 1) to easily view the sample-level annotations in a GEO series and/or 2) to perform custom tidying operations on a GEO series and export the data to other tools. By facilitating these processes, TidyGEO promises to accelerate the research process and reduce barriers for biologists.

## Methods

*TidyGEO* is a Shiny web application (http://shiny.rstudio.com) hosted in a Docker container (https://docker.com)^42^. The user begins by entering a GEO Series identifier. Initially, TidyGEO displays metadata about the series, including organism, experiment type, summary, and publication information. A link to the series on the GEO website is also provided. If the user chooses to “Import” the series, TidyGEO uses the *GEOquery* R package^28^ to download sample-level annotations, assay data, and platform annotations. Options for manipulating the data are organized based on high-level operations: 1) tidying sample-level annotations (“clinical” data), 2) tidying assay data, 3) tidying platform annotations (feature data), and 4) merging data. These sub-pages can be accessed in any order, although TidyGEO attempts to guide the user through a logical sequence of steps.

### Clinical data

Clinical data (sample-level annotations) describe the research subjects (samples) from which assay measurements were derived, providing insight into factors that may explain biological variation. Typically, scientists use clinical variables either as the primary variable of interest or as covariates. These variables may also be used to identify sample subsets that meet certain criteria—for example, patients with a certain blood type.

TidyGEO provides functionality for manipulating clinical data so that they can more easily be used in downstream analyses. This functionality is presented to the user in a series of tabs that allows users to 1) select informative columns, 2) shift cells due to missing data, 3) split key-value pairs, 4) split columns based on delimiters, 5) rename columns, 6) substitute values in a given column, 7) filter samples, and 8) save the tidied data to a file. TidyGEO guides users from tab to tab via “Next” and “Back” buttons; however, users need only perform tasks that are relevant to a particular dataset.

As users visit the tabs and manipulate data, views of the data are displayed on the right panel. The default option is a tabular view (data represented as rows and columns). Users can navigate the data horizontally or vertically using control buttons. A second option provides graphical summaries of the data. Users select a column to visualize. If the variable is numeric, TidyGEO generates a histogram and allows the user to select the bin size. If the variable is categorical, TidyGEO generates a bar plot. In either case, the user can download the figure (PDF, JPG, or PNG format) and specify the color used, image width and height, and font size. The graphs are generated via the ggplot2 package^43^.

The tabs that support tidying clinical data are described below.

#### Selecting informative columns

Many sample-level annotations provide information that is not useful for quantitative analyses. TidyGEO users can select individual columns to be removed. Alternatively, they can indicate that they wish to remove all columns for which 1) all samples have the same value, 2) all samples have unique values, 3) all values are dates, or 4) all values are Web addresses. TidyGEO then removes these columns.

#### Shifting cells

When an annotation is missing for a given sample, data values for that sample may be shifted leftward to other columns. TidyGEO users can shift such values to the right so that data for each variable is stored in a single column across all samples. Users indicate which columns contain relevant values and then specify a pattern that indicates which cells have relevant data. TidyGEO then extracts these values to a new column.

#### Splitting key-value pairs

When annotations are represented as key-value pairs (e.g., “treatment:control” or “sex=female”) in a given column—and the key is identical for all samples—users can indicate that the key should be used as the column name and the column should contain the respective values. Users indicate the delimiter that separates the key-value pairs (e.g., “:” or “=”). For more complicated scenarios, a regular expression can be used.

#### Splitting columns

When multiple data points (or key-value pairs) are nested within cells in a given column, users can split those columns by specifying a delimiter (or regular expression). TidyGEO then separates these values into multiple columns.

#### Renaming columns

Users can rename a column to better reflect its semantics and to improve consistency of column names.

#### Substituting column values

Users can replace all instances of a given value with an alternative value. For example, if “F” were used to represent females, the user could change these values to “Female.” Regular expressions can be used for more advanced scenarios.

#### Filtering samples

For a given column, TidyGEO users can indicate that all samples with a given value should be excluded. For example, all males could be excluded.

#### Saving the data

Users can download the data in any of five file formats: comma-separated value (CSV), tab-separated value (TSV), JavaScript Object Notation (JSON), Microsoft Excel, or the format required by BRB-ArrayTools. BRB-ArrayTools is a spreadsheet-based system for analyzing gene-expression microarray, copy-number, methylation and RNA-Seq data^38^.

### Assay data

Researchers may wish to summarize assay measurements. For example, Affymetrix gene-expression microarrays map measurements to “probe sets,” which quantify expression for gene regions^44^. In many cases, multiple probe sets are associated with a single gene. Thus, probe-set values may be difficult to interpret for researchers who are interested in gene-level measurements. Alternatively, for RNA-Sequencing data, values may be quantified at the transcript level. Again, researchers may wish to summarize the data at the gene level. In its “Assay data” panel, TidyGEO enables users to examine the platform annotations for a given series and select an alternative way of summarizing the abundance measurements. For example, if Affymetrix data had been summarized at the probe-set level and the platform annotations provided a gene symbol (or other gene identifier) for each probe set, the user could summarize the measurements at the gene level. In another example, if the platform annotations indicated the cytogenetic band associated with each measurement, the user could summarize the data based on cytogenetic band. When summarizing, the user can select from the following options: mean, median, minimum, or maximum. Alternatively, the “keep all” option assigns the specified identifiers to the abundance measurements but does not collapse the values.

Regardless of the summarization method used, TidyGEO enables researchers to filter the assay data based on values in a given column. For example, a researcher might retain only rows in which the measurement was higher than a given threshold. As with the clinical data, the user can create visualizations of each column in the assay data and download these graphics.

In GEO, clinical annotations for each sample are stored on a row-by-row basis, whereas assay measurements for each sample are stored on a column-by-column basis. To support easier merging of these data types, TidyGEO enables users to transpose the assay data with the click of a button. In addition, the user can export the assay data in a variety of formats.

### Feature data

Platform annotations in GEO suffer from many of the same limitations as sample-level annotations. For example, in some cases, extraneous columns are present, or multiple identifiers are stored in the same cell. In other cases, data are stored as key/value pairs, and values can become shifted when data are missing. To address these problems, TidyGEO provides options for tidying platform annotations: selecting informative columns, shifting empty cells, splitting key/value pairs, and splitting columns based on a delimiter. This process is independent of the process of analyzing assay data, but it can help with the latter. Tidying platform annotations can ensure that assay data are summarized using consistent and relevant identifiers, which is particularly important for deriving inferences that span multiple datasets and mapping data to external resources, such as pathway databases^45^.

As with the other panels in TidyGEO, users can create visualizations of the platform annotations in each column, download graphics, and download the platform data in a variety of formats.

### Merging data

Many researchers will wish to identify relationships between sample-level annotations and assay measurements. TidyGEO enables users to merge these data types and export them as combined data files. If a user has used other panels in TidyGEO to filter data, the user can choose whether to perform an inner join (retain only samples that are present for both data types), left or right join (retain all samples from the first or second data type, respectively), or a full join (retain data from samples in either dataset).

### Supporting computational reproducibility

When an analysis is computationally reproducible, the researcher who performed the analysis maintains a detailed record of the analysis steps that were performed and is able to repeat those steps, leading to the same (or similar) results^46^. However, this process is typically tedious when the analysis was performed using a graphical interface because the researcher must manually record each step in detail. An additional complication is that the user interface may change as new versions of the software are released, so these descriptions may become obsolete. TidyGEO addresses this problem by generating reproducible scripts. As a user processes a GEO series, TidyGEO records the code that was used to perform those tasks. On each panel, TidyGEO provides an option for the researcher to download an R script that contains the code, along with comments that explain the tasks being performed.

## Declarations

### Ethics approval and consent to participate

Brigham Young University’s Institutional Review Board approved this study under exemption status. TidyGEO uses data collected from a public repository. We play no part in patient recruiting or in obtaining consent.

### Consent for publication

Not applicable.

### Availability of data and materials

Source code for TidyGEO is available at https://github.com/srp33/TidyGEO; this repository also includes a developer’s guide to provide insight into how the application works. Packaged within a software-container image, TidyGEO can be executed on a researcher’s own computer; we also provide a public implementation for demonstration purposes (https://bioapps.byu.edu/TidyGEO).

## Authors’ contributions

AM conceived the project, developed the software, collected data, analyzed data, created visualizations, and wrote the manuscript. AS developed the software. BIQ developed the software. GS collected data. SRP conceived the project, supervised the project, provided computational resources, collected data, developed the software, wrote the manuscript, and edited the manuscript.

## Acknowledgements

We thank James Wengler, Teancum Paquette, and Ifeanyichukwu Nwosu for providing helpful feedback on *TidyGEO*’s usability.

